# RasGRP1 (CalDAG-GEF-II) Mediates L-DOPA-induced Dyskinesia in a Mouse Model of Parkinson Disease

**DOI:** 10.1101/739631

**Authors:** Mehdi Ishragi, Uri Nimrod Ramirez Jarquin, Neelam Shahani, Supriya Swarnkar, Nicole Galli, Oscar Rivera, George Tsaprailis, Catherina Scharager-Tapia, Gogce Crynen, Alessandro Usiello, Srinivasa Subramaniam

## Abstract

The therapeutic benefits of L–3,4–dihydroxyphenylalanine (L-DOPA) in Parkinson disease (PD) patients diminishes with the onset of abnormal involuntary movements (L-DOPA induced dyskinesia), a debilitating motor side effect. L-DOPA induced dyskinesia are due to altered dopaminergic signaling in the striatum, a brain region that controls motor and cognitive functions. However, the molecular mechanisms that promote L-DOPA-induced dyskinesia remain unclear. Here, we have reported that RasGRP1 (also known as CalDAG-GEF-II) physiologically mediated L-DOPA induced dyskinesia in a 6-hydroxy dopamine (6-OHDA) lesioned mouse model of PD. In this study, L-DOPA treatment rapidly upregulated RasGRP1 in the striatum. Our findings showed that RasGRP1 deleted mice (*RasGRP1^−/−^*) had drastically diminished L-DOPA-induced dyskinesia, and *RasGRP1^−/−^* mice did not interfere with the therapeutic benefits of L-DOPA. In terms of its mechanism, RasGRP1 mediates L-DOPA-induced extracellular regulated kinase (ERK), the mammalian target of rapamycin kinase (mTOR) and the cAMP/PKA pathway and binds directly with Ras-homolog-enriched in the brain (Rheb), which is a potent activator of mTOR, both in vitro and in the intact striatum. High-resolution tandem mass tag mass spectrometry analysis of striatal tissue revealed significant targets, such as phosphodiesterase (Pde1c), Pde2a, catechol-o-methyltransferase (comt), and glutamate decarboxylase 1 and 2 (Gad1 and Gad2), which are downstream regulators of RasGRP1 and are linked to L-DOPA-induced dyskinesia vulnerability. Collectively, the findings of this study demonstrated that RasGRP1 is a major regulator of L-DOPA-induced dyskinesia in the striatum. Drugs or gene-depletion strategies targeting RasGRP1 may offer novel therapeutic opportunities for preventing L-DOPA-induced dyskinesia in PD patients.

## INTRODUCTION

The loss of substantia nigral projections neurons, which results in decreased dopamine levels in the striatum, is the principle cause of PD. A drug known as L-DOPA, a precursor of dopamine synthesis, effectively alleviates motor stiffness in PD, but its therapeutic benefits are markedly limited by its debilitating dyskinetic side effects. Previous studies have shown that L-DOPA-induced dyskinesia is mediated by the activation of ERK and mTORC1 signaling in the striatum^1–3^. The inhibitors of these signaling may prevent L-DOPA-induced dyskinesia without affecting the beneficial effects of L-DOPA on limb motion^3, 4^. Ras-GRF1, which is abundant in the cortex, hippocampus, and striatum, is also known to activate ERK in the striatum and regulate L-DOPA-induced dyskinesia^5^. We found that Rhes, a striatal-enriched GTPase/SUMO-E3-like protein, binds and activates mTORC1 signaling and promotes L-DOPA-induced dyskinesia^6^, consistent with those of an independent report^7^. However, the striatal regulators that mediate both ERK and mTORC1 signaling and L-DOPA-induced dyskinesia remain unknown.

RasGRP1, a guanine-nucleotide exchange factor (GEF) for H-Ras that signals ERK, is highly expressed in hematopoietic cells and is regulated by calcium and diacylglycerol^8–11^. RasGRP1 is known to play a role in T- and B-cell proliferation and has been implicated in leukemia and lupus^11–15^. RasGRP1 is enriched in selected brain regions that control motor and cognitive functions, including the striatum^10^. Despite its high expression in brain regions, its role in neuronal functions remains unclear. Previously, we found that RasGRP1 interacts and acts as a GEF for Rhes and promotes amphetamine-induced hyperactivity via the striatal protein-protein complex known as Rhesactome^16^. Here, we have reported a major role for RasGRP1 in L-DOPA-induced dyskinesia and demonstrated that RasGRP1 is causally linked to L-DOPA-induced dyskinesia and robustly activates ERK and mTORC1 pathways in the striatum. Using high-end quantitative proteomic analysis of WT and *RasGRP1^−/−^* mice, we have identified multiple striatal targets downstream of RasGRP1 activation that may be involved in L-DOPA-induced dyskinesia.

## RESULTS

### RasGRP1 promoted L-DOPA-induced dyskinesia in a mouse model of Parkinson disease

We hypothesized that RasGRP1 may be an upstream regulator of L-DOPA-induced dyskinesia due to the following reasons: a) L-DOPA treatment of mice with unilateral 6-hydroxydopamine (6-OHDA) lesions of the nigrostriatal pathway augmented striatal ERK and mTOR signaling^1–3^; b) Rhes, a striatal-enriched protein that activates mTOR, is involved in L-DOPA-induced dyskinesia^6, 17, 18^; and c) RasGRP1 regulated the synaptic localization of Rhes and RasGRP1, and Rhes co-expression strongly activated both ERK and mTORC1 signaling in a cell culture^16^. To test our hypothesis, we subjected WT and *RasGRP1^-/-^* (RasGRP1 KO) mice to a well-established 6-OHDA lesion model of L-DOPA-induced dyskinesia as in our earlier study^6^. Fig 1A shows the timeline of the 6-OHDA lesion and L-DOPA-induced dyskinesia analysis. We observed 6-OHDA-induced PD-like symptoms in the drag test, rotarod, and turning test, which were similar between WT and RasGRP1 KO mice (Fig. 1B). The open field test did not show obvious differences (Fig. 1B). Daily treatment of unilaterally-6-OHDA lesioned mice with 5 mg/kg L-DOPA produced dyskinesia (abnormal involuntary movements, AIMs, measured after for 1, 4, 7, 11, 14, and 17 days) in WT mice, which was robustly diminished in RasGRP1 KO mice (Fig. 1C). We observed in WT mice that L-DOPA induced the progressive development of severely abnormal axial, limb, and locomotor movements, which were dramatically attenuated in RasGRP1 KO mice (Fig. 1D). The dyskinetic influences of L-DOPA increased over the course of time in WT mice (days 1 - 17), but the increase was much less in RasGRP1 KO mice between 20 and 80 min after L-DOPA injection, though this was followed by a marked decline two hours later. The rate of decline was similar in WT and RasGRP1 KO mice, suggesting that RasGRP1 did not alter L-DOPA turnover (Fig. 1E). These observations have shown that RasGRP1 is a critical regulator of L-DOPA-induced dyskinesia.

**Figure 1.**
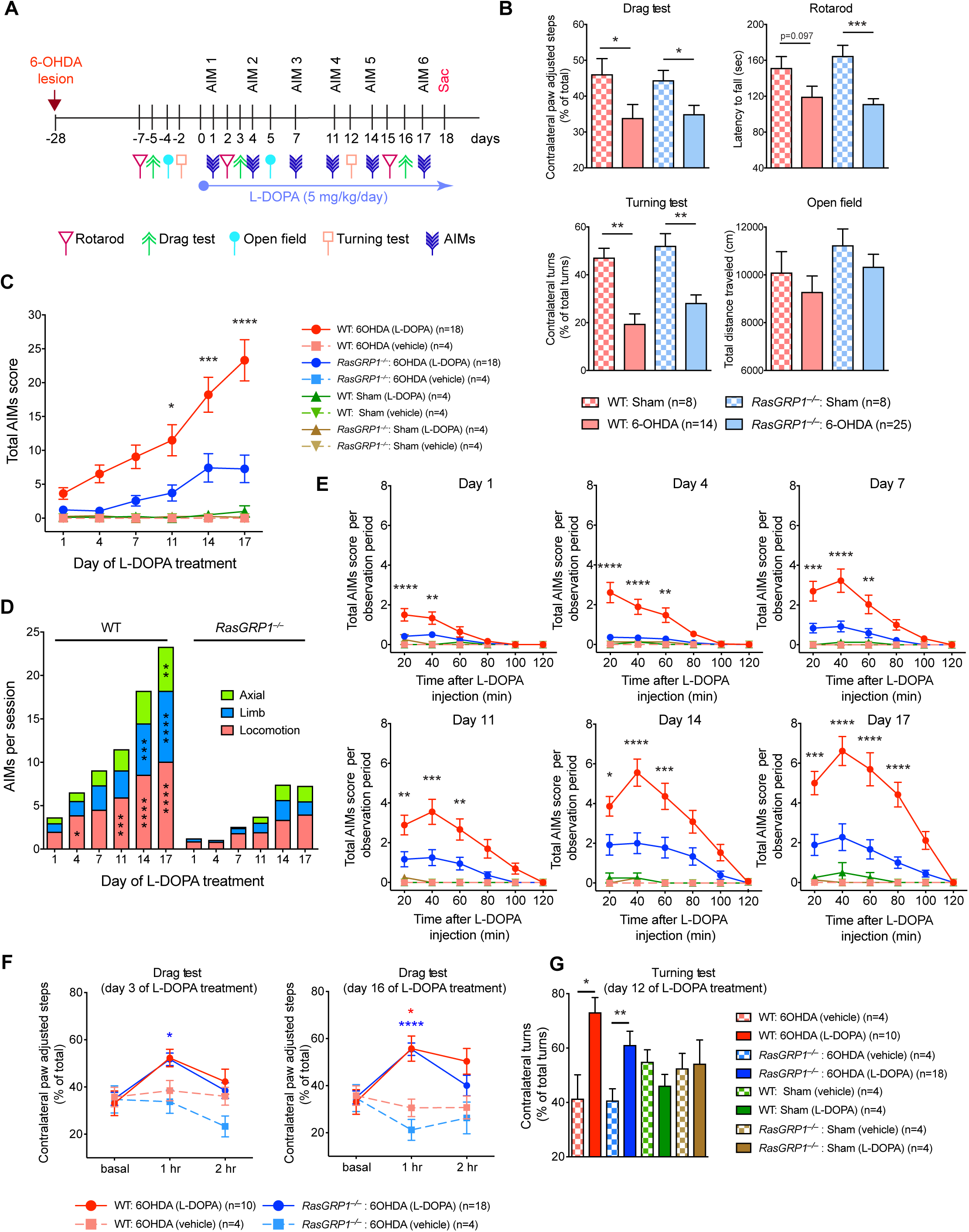
RasGRP1 promotes L-DOPA induced dyskinesia. (A) L-DOPA-induced dyskinesia scheme. (B) Shows drag test, rotarod, turning test and open field test, for the indicated genotypes for sham or 6-OHDA-lesioned mice. Total AIM scores (C) or AIMs per session (D) (Axial, limb, or locomotion) for the indicated 6-OHDA-lesioned WT and RasGRP1 KO mice, vehicle or L-DOPA injected mice. (E) Shows total AIMs score per observed period (Day 1-Day 17), after injection of L-DOPA. (F) Shows drag test on day 3 and day 16, (G) turning test on day 12 after L-DOPA injection (n = 4-18/group, One-way ANOVA followed by Bonferroni post hoc test, repeated measures Two-Way ANOVA followed by Bonferroni post hoc test, *p<0.05; **p<0.01; ***p<0.001; **** p<0.0001).

### RasGRP1 deletion did not influence the anti-Parkinson effects of L-DOPA

Next, we investigated whether RasGRP1 KO had any influence on the anti-Parkinson effect of L-DOPA. We found that in both WT and RasGRP1 KO-lesioned mice, administering L-DOPA decreased Parkinson symptoms as measured by the drag test (on day 3 and day 16, Fig. 1F) and the turning test (day 12, Fig. 1G). As expected, sham injections produced no defects in the drag test (Supplementary Fig. 1). The open field or rotarod test was also used as an anti-Parkinson disease readout, but we did not see any difference in total distance travelled or rotarod test results between the genotypes and sham treatment (Supplementary Figs. 2 and 3). Thus, RasGRP1 promoted the adverse effects of L-DOPA but not its therapeutic motor effects. Moreover, RasGRP1 KO mice displayed no significant changes in basal motor behavior or coordination^16^, as shown in Supplementary Figs 3–5, which would not likely contribute much to the altered L-DOPA responses we observed.

### RasGRP1 was upregulated during L-DOPA-induced dyskinesia and promoted ERK, mTOR, and GluR1 signaling in the striatum of dyskinetic mice

Research has shown that dopamine 1 (D1) and dopamine 2 (D2) receptors of medium spiny neurons (MSN) in the striatum may play different roles in L–DOPA dyskinesia. For example, Santini et al. showed that ERK and mTOR signaling selectively enhanced L-DOPA treatment in D1 containing neurons^3^. Consistent with these data, the stimulation of mTOR signaling by L-DOPA has been shown to be abolished by D1 antagonists and unaffected by D2 antagonists^3^. Similarly, we found that Rhes, which is predominantly expressed in D1 and D2 MSN, promoted mTOR in L-DOPA-induced dyskinesia^6^. As RasGRP1 KO mice showed diminished dyskinesia, we tested WT and RasGRP1-KO striatal tissue on L-DOPA-induced ERK, mTOR, and other signaling in the striatum. We observed several findings related to signaling in our model. First, we confirmed that lesions produced more than a 60% loss of tyrosine hydroxylase (TH)^+^ in both WT and RasGRP1 KO mice (Fig. 2A, B), which indicated the onset of PD-like symptoms. Intriguingly, we found that RasGRP1 levels were upregulated after L-DOPA injection, consistent with a previous report in a rat model of L-DOPA-induced dyskinesia^19^ (Fig. 2A, B). This increase was dependent on L-DOPA administration, as we found no increase of RasGRP1 in 6-OHDA lesion vehicle control (Supplementary Fig. 4). We measured mTORC1 activity according to levels of phosphorylation of ribosomal protein S6 kinase (S6K) at T389 and phosphorylation of S6 at S235/236 in vivo and phosphorylation of eukaryotic translation initiation factor 4E (eIF4E)-binding protein 1 (4EBP1) (T37/46) at a site that primes p4EBP1 for subsequent activity phosphorylation at S65. We were unable to detect p4EBP1 S65, as the antibodies for this site did not work for brain lysate. We also found that mTORC2 activity, as measured by phosphorylation of Akt (S473), was upregulated in 6-OHDA-lesioned WT mice but not in RasGRP1 KO mice (Fig. 2A, B). Previous findings have shown that Akt S473 phosphorylation was upregulated in a monkey model of L-DOPA-induced dyskinesia^20^. In addition, pGlur1 S845, pERK (T202/Y204), and the PI3K target, pAkt (T308), were also highly upregulated in WT but not RasGRP1 KO mice (Fig. 2A, B). We found that Rheb and Rhes levels, were significantly downregulated in the lesion side of RasGRP1 KO mice (Fig. 2, A, B), consistent with our earlier report^16^, indicating that RasGRP1 may physiologically stabilize these proteins in the striatum. Together, these biochemical studies indicated that a) RasGRP1 is upregulated in an L-DOPA-dependent manner; b) RasGRP1 KO prevents the upregulation of L-DOPA-induced mTOR, ERK, Akt, Glur1 signaling in the striatum; and c) Rhes and Rheb, activators of mTOR, are downregulated in the striatum of RasGRP1 KO mice compared to WT mice during L-DOPA-induced dyskinesia.

**Figure 2.**
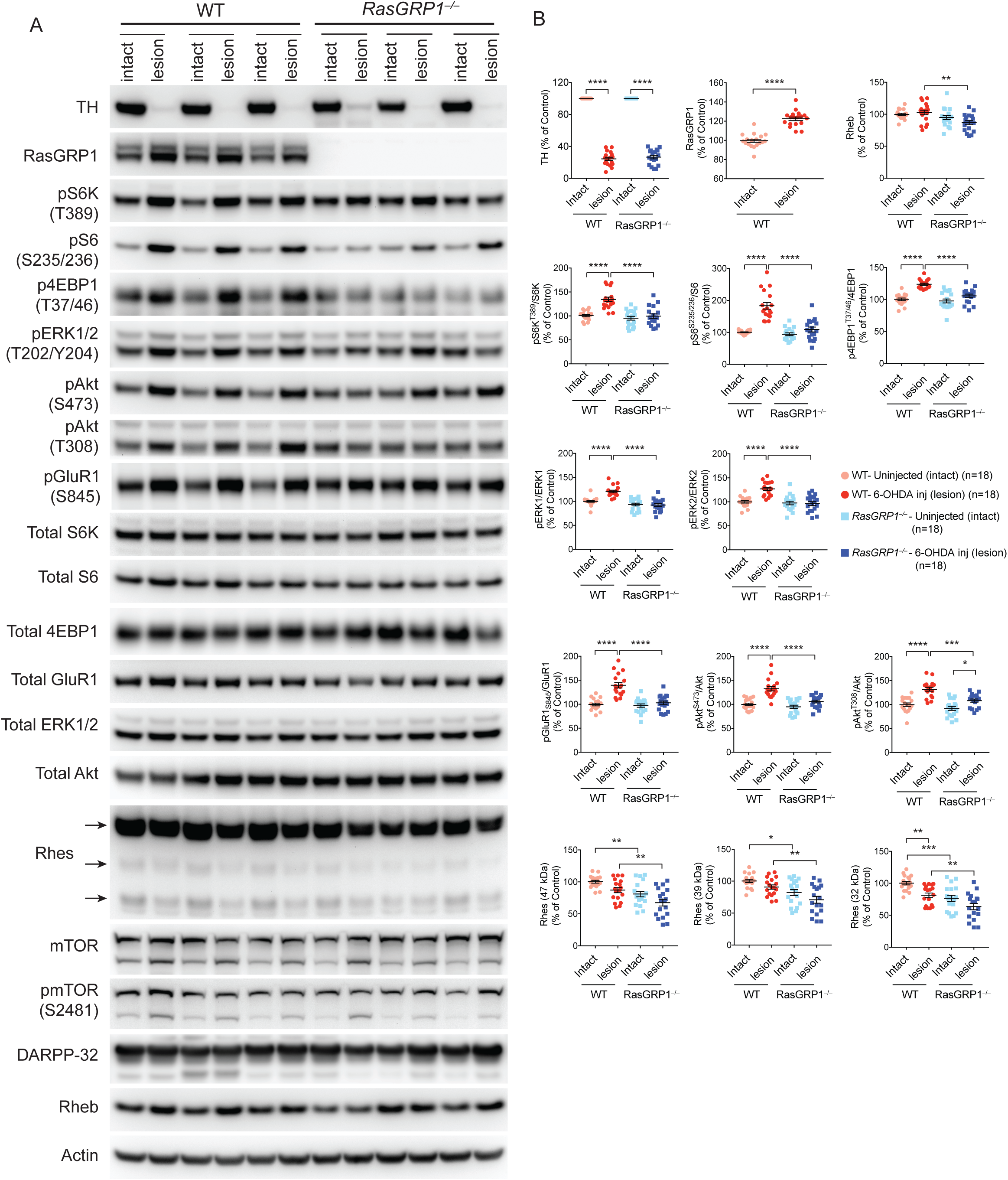
RasGRP1 mediates L-DOPA induced mTOR and ERK signaling in the striatum. (A) Western blot analysis of intact and 6-OHDA-lesioned striatum of WT and RasGRP1 KO mice. (B) Quantification of the indicated signaling In WT or RasGRP1-KO intact side or lesioned side of the striatum (n = 18/group, One-way ANOVA followed by Tukey’s multiple comparison test, *p<0.05; **p<0.01; ***p<0.001; ****p<0.0001).

### RasGRP1 mediated mTOR signaling in a PI3K-dependent and ERK-independent manner

Next, we examined the potential mechanisms by which RasGRP1 upregulation in L-DOPA induced dyskinesia promoted mTOR and ERK signaling. To do this, first, we transiently overexpressed RasGRP1 in HEK293 cells, which were transfected with a control vector (His) or His-RasGRP1 cDNA. Then, we incubated HEK293 cells in serum-free F12 medium with full amino acids (plus AA) or F12 medium that lacked L-leucine (minus AA), which is a potent inducer of mTORC1^21^. After 2 h, cells in minus-AA medium were re-stimulated with L-leucine (3 mM). In full-AA conditions, compared to control cells, RasGRP1-expressing cells had twice to three times as much mTORC1 activity as measured by the phosphorylation of S6K at T389 (pS6K-T389) and p4EBP1-S65 (Figs. 3 A, B). In minus-AA medium, mTORC1 activity was 50% lower in RasGRP1 expressing cells yet was higher than the control. Upon re-stimulation with L-leucine for 15 min, mTORC1 activity rapidly returned to levels comparable with those in full-AA conditions. This indicated that RasGRP1 mediated AA-mediated mTORC1 activity. RasGRP1 expression induced the constitutive phosphorylation of ERK, which is not sensitive to amino acids (Fig. 3C).

**Figure. 3.**
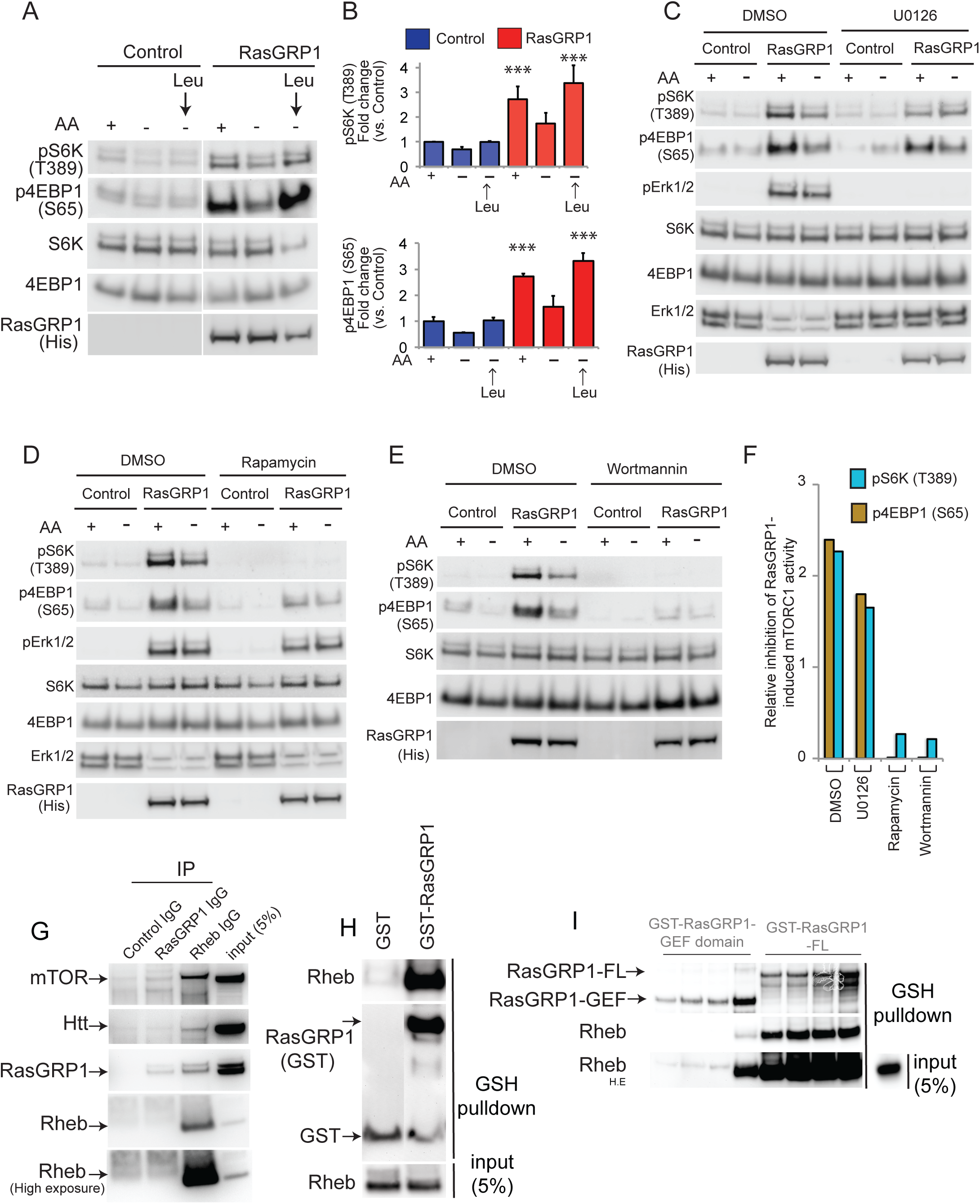
RasGRP1 regulates mTOR activation, independent of ERK signaling. (A) RasGRP1 mediates AA-induced mTORC1 activity. HEK293 cells (grown in DMEM + serum) were transfected with His (control) or His-RasGRP1constructs (0.5µg each), after 36-48 hr, the cells were exposed to serum free media (F12+) containing all amino acids (AA+) or serum free media (F12–) that lacks L-Leucine (AA–) for 2 hr, and wherever indicated F12– media was stimulated with L-Leucine (3 mM) for 15 min. Cell lysates were probed for p-S6K (T389), p-4EBP1 (S65) and other indicated proteins by Western blotting. (B) Displays quantification of A. ***p < 0.001 vs. control, Student’s-t-test. (C) RasGRP1-mediated mTORC1 activity is independent of ERK signaling. HEK293 cells were grown in A, replaced with AA+ or AA– media with DMSO (0.01%) or U0126 (10 µM) for 2 hr. Cell lysates were prepared and probed using Western blotting for indicated protein. (D) Rapamycin abrogates RasGRP1-mediated mTORC1 signaling. Cells were transfected as in A followed by changing the medium to AA+ or AA– as in C with DMSO or rapamycin (500 nM) and probed for indicated protein by Western blotting. (E) Wortmannin abrogates RasGRP1-mediated mTORC1 activity. Cells were transfected as in A, and the AA+ and AA– media was treated with DMSO or wortmannin (100 nM) for 2 hr, followed by detection of indicated protein through Western blotting. (F) Indicates relative inhibitory potency of different inhibitors, on the RasGRP1-mediated mTORC1 activity. (G). Western blot showing in vitro Rheb and RasGRP1 binding in the striatum, in vivo. (H). Western blot showing recombinant Rheb and RasGRP1 protein interaction in vitro. (I). Western blot showing GEF domain and RasGRP1-FL domain interaction with Rheb in vitro.

Next, we tested the effects of ERK inhibition on RasGRP1-mediated AA-mTORC1 signaling. While U0126, a potent inhibitor of mitogen-activated protein kinase (MEK) abrogated ERK signaling, it had negligible effects on RasGRP1-induced pS6K (Thr389) and p4EBP1 (Ser65, Fig. 3C). However, rapamycin, the mTORC1 inhibitor, which did not alter RasGRP1-induced ERK signaling, markedly attenuated RasGRP1-induced AA-mTORC1 signaling (Fig. 3D). Like rapamycin, the PI3K inhibitor wortmannin abolished RasGRP1-induced mTORC1 signaling (Fig. 3E). Collectively, rapamycin and wortmannin but not U0126 blocked RasGRP1-induced mTORC1 activation (Fig. 3F). The depletion of endogenous RasGRP1 using short hairpin RNA (shRNA) in HEK cells also diminished mTORC1 signaling (Supplementary Fig. 5). Together, this data suggested that RasGRP1 physiologically activates mTORC1 signaling by regulating catalytically important residues on pS6K and p4EBP1 via a PI3K-sensitive pathway. In addition, using a pharmacological approach, we found no significant cross talk between ERK and mTORC1 signaling induced by RasGRP1.

### RasGRP1 directly interacted with Rheb in the striatum

As there was no cross talk, we predicted that RasGRP1 would activate ERK and mTORC1 signaling in two independent and parallel pathways. Research has shown that RasGRP1 can activate ERK via GTPase H-Ras^8, 10^, but we wondered how RasGRP1 activates AA-induced mTORC1. As Rheb, which directly binds and activates mTOR, mediates AA-induced mTORC1 activity^22^, we hypothesized that RasGRP1 may promote mTORC1 activity by interacting with Rheb in the brain. To test this, we co-immunoprecipitated Rheb and RasGRP1 from the brain’s striatum. As hypothesized, Fig 3G shows that Rheb effectively co-immunoprecipitated with RasGRP1 and mTOR, a known Rheb interactor. Rheb also coprecipitated huntingtin, which was consistent with our previous report^23^. However, the RasGRP1 antibody appeared inefficient for immunoprecipitation, as it only moderately enriched RasGRP1 (Fig. 3G). Next, we investigated whether RasGRP1 interacted directly with Rheb in vitro. Co-incubation of bacterially purified GST-FL-RasGRP1 and Rheb revealed their robust interaction (Fig. 3H). As shown in Fig. 3I, the interaction appeared strong with FL-RasGRP1 compared to the RasGRP1 GEF domain (1-470 amino acids,^11^. Thus, for the first time, we showed that RasGRP1 can directly interact with Rheb in the striatum. Collectively, this data suggested that RasGRP1 may promote mTORC1 activity in the brain via Rheb GTPase.

### Quantitative striatal proteomic analysis of WT and *RasGRP1^−/−^* mice revealed novel RasGRP1 targets with potential involvement in L-DOPA-induced dyskinesia

To understand the mechanisms by which RasGRP1 might elicit L-DOPA-induced dyskinesia, we undertook quantitative and comparative proteomics profiling of WT and RasGRP1 KO striatum with high-resolution mass spectrometry coupled to liquid chromatography (LC-MS/MS) based on tandem mass tags (TMT) designed for phosphoprotein enrichment. We isolated the striatum 20 min after L-DOPA administration from three WT mice that showed severe dyskinesia and three RasGRP1 KO mice that showed no dyskinesia in response to L-DOPA in a PD model (Fig. 4A). We labelled each lysate with TMT labelled six-plex reagents (ThermoFisher) as indicated in Table 1.

**Table 1.**

| Table. 1 | WT-intact (A) | WT-lesion (B) | *RasGRP1*-KO intact (C) | *RasGRP1*-KO lesion (D) |
| --- | --- | --- | --- | --- |
| Plex 1 label | 126 (Kit1) | 127 (Kit1) | 128 (Kit1) | 129 (Kit1) |
| Plex 2 label | 128 (Kit2) | 129 (Kit2) | 130 (Kit2) | 131 (Kit2) |
| Plex 3 label | 126 (Kit2) | 127(Kit2) | 130 (Kit1) | 131 (Kit1) |

**Figure 4.**
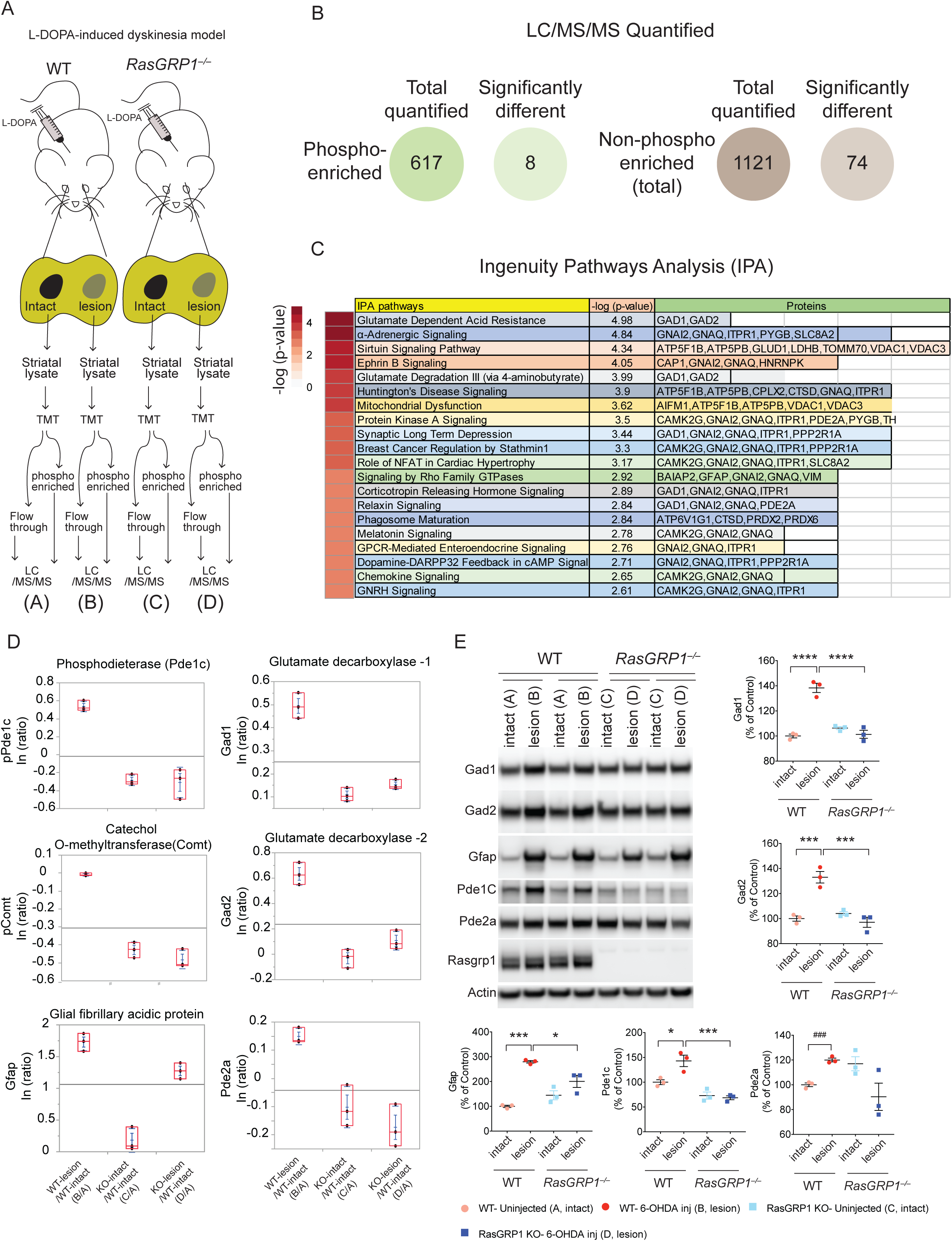
Quantitative proteomics of the striatum of WT and RasGRP1-KO dyskinesia animals. (A) Indicates scheme of isolation of striatal tissue from animals that were subjected to L-DOPA induced dyskinesia, followed by LC-MS/MS. (B) Depicts total number of quantifiable proteins that are enriched for phosphorylated and non-phosphorylated forms. (C) Indicates IPA analysis for significantly altered proteins. (D) Indicates quantification (ratio) of selected examples of LC/MS/MS of proteins in comparison to intact WT striatum. (E) Shows western blotting and quantification of indicated proteins from the intact and lesioned striatum of WT and RasGRP1-KO. (n = 3/group, One-way ANOVA followed by Tukey’s multiple comparison test. *p<0.05; ***p<0.001; ****p<0.0001) or Student’s t-test. ^###^p<0.001.

Loading bias was minimal and was removed by normalizing it with the total peptide amount. We quantified 617 phosphorylated proteins in all groups. ANOVA results indicated eight proteins were significantly regulated between the comparison groups (B/A, C/A, and D/A) (FDR 0.10) and then Turkeys HSD was used as post hoc test to find out which pairs were different from each other (alpha = 0.05) (Fig. 4B, data files S1, S2, S3). Similarly, ANOVA results indicated seventy four proteins were significantly regulated, out of 1121 proteins identified, between the comparison groups (B/A, C/A, and D/A) (FDR 0.10) and then Turkeys HSD was used as post hoc test to find out which pairs were different from each other (alpha = 0.05, Fig. 4B, data files S4, S5, S6). Ingenuity Pathway Analysis (IPA) revealed that signaling pathways related to glutamate-dependent acid resistance, *α*-adrenergic signaling, sirtuin signaling, ephrin signaling, and glutamate degradation, Huntington disease pathways, mitochondrial dysfunction targets, protein kinase A signaling, and others were highly significantly upregulated in WT but not in RasGRP1 KO striatum (Fig. 4C, data file S7). As shown in Fig. 4D, selected phosphorylated and total proteins upregulated in the lesion side of WT but not in RasGRP1 KO striatum. We observed that the phospho-enrichment of phosphodiesterase 1c (Pde1c), total Pde2a, both with a dual-specificity for the second messengers, cAMP and cGMP^24, 25^, was higher in WT than RasGRP1 KO striatum. This had demonstrable relevance in L-DOPA-induced dyskinesia, because the cAMP pathway^26–28^ and PDE activation have been linked to L-DOPA-induced dyskinesia in a mouse and monkey model of PD^29, 30^. The specific role of Pde1c and Pde2a in regulating L-DOPA-induced dyskinesia is unknown. Likewise, the phosphorylation of Comt, an enzyme that catalyzes the degradation of catecholamines (including the neurotransmitters L-DOPA, dopamine, and epinephrine) is upregulated in WT but not in RasGRP1 KO mice. Inhibitors of Comt can enhance L-DOPA-induced dyskinesia in monkeys^31^, indicating that the upregulation of Comt is an adaptative response designed to degrade L-DOPA, a feature that is absent in RasGRP1 KO mice. GFAP upregulation is found in 6-OHDA rodent model of PD^32, 33^. Similarly, we found a dramatic upregulation of GFAP in the lesion side of both WT and RasGRP KO, although it is slightly less in the KO striatum (Fig. 4D). Gad 1, also known as Gad 67 and Gad 2, also known as Gad 65, catalyze the production of gamma-aminobutyric acid (GABA) and are upregulated in WT but not in RasGRP1 KO striatum (Fig. 4D). The loss of Gad 67 has diminished L-DOPA-induced dyskinesia in mouse models of PD^34^. We used Western blotting with a tested antibody to validate the upregulation of Gad 1, Gad 2, Gfap, Pde1c and Pde2a observed in quantitative LC/MS/MS in the lesioned area of WT mice but not in RasGRP1 KO striatum (Fig. 4E). However, not all proteins were downregulated in RasGRP1 KO mice. For example, complexin-1, visinin-like protein, glutamate dehydrogenase 1, guanine nucleotide-binding protein G (g) subunit alpha, and lactate dehydrogenase were upregulated in RasGRP1 KO mice but not WT mice (data file S6). Collectively, LC/MS/MS data indicated that RasGRP1 acts upstream in response to L-DOPA and regulates a specific but diverse set of proteins to promote L-DOPA-induced dyskinesia. This notion is strengthened by the fact that some of these proteins have been implicated in L-DOPA-induced dyskinesia in independent studies^29–31^.

## DISCUSSION

After 10 years of treatment, more than 95% of PD patients who receive L-DOPA medication develop dyskinesia, an uncontrolled motor deficit for which no effective therapy exists. There is an immediate need to identify novel therapeutic targets for L-DOPA-induced dyskinesia. Our study demonstrated that RasGRP1 is a major therapeutic target, because our data indicated that a) RasGRP1 is induced upon L-DOPA administration, b) RasGRP1 is causally linked to L-DOPA-induced dyskinesia, and c) RasGRP1 physiologically mediates L-DOPA-induced activation of ERK and mTOR, which are linked to L-DOPA-induced dyskinesia. RasGRP1 is a better target, because although mTOR inhibitors, such as rapamycin, or ERK inhibitors, such as U0126, have been shown to prevent L-DOPA-induced dyskinesia in mouse models^3, 4, 7^, these drugs are strong inhibitors of protein synthesis and have associated toxicity. In addition, they broadly inhibit targets in unwanted regions. Further, RasGRP1 KO mice have no gross phenotype, and they are fertile and have no significant changes in the basal motor activity^16^, Supplementary Figs. 1-3). RasGRP1 KO mice does show mild defects (20%) in thymocyte development^35^. Thus, drugs blocking RasGRP1 or shRNA approaches to deplete RasGRP1 in adults may produce fewer adverse effects and strong beneficial effects in preventing L-DOPA-induced dyskinesia.

Our study showed that RasGRP1 induces two major pathways, ERK and mTOR signaling, in the striatum during L-DOPA-induced dyskinesia. Cell culture data demonstrated that there was no cross talk between ERK and mTORC1 signaling and suggested a role for PI3K in these interactions. We proposed a model to show how RasGRP1 might activate both ERK and mTOR pathways during L-DOPA-induced dyskinesia (Fig. 5). We predicted that RasGRP1 would form complexes with more than one small GTPase in the striatum during L-DOPA-induced dyskinesia (Fig. 5). For example, upregulated RasGRP1 can form two kinds of GEF complex, a RasGRP1-Ras complex and a RasGRP1-Rheb complex, to activate ERK and mTOR signaling, respectively. This notion was consistent with the actions of other GEFs that activate multiple small GTPases^36, 37^. We predicted that upregulated RasGRP1 would act like a “master GEF” for H-Ras, Rhes, and Rheb, deviating from the conventional notion that it acts as one GEF for one small GTPase. Such “GEF-GTPasome” complexes may afford the flexibility needed to develop the abnormal involuntary movements observed in L-DOPA-induced dyskinesia. Supporting the notion of the existence of GEF-GTPasome we found that Rheb and Rhes both interacted directly with RasGRP1 in vitro (data not shown). We also predicted that RasGRP1 would be upregulated in response to dopamine D1 receptor super sensitivity in PD. To support this notion, we found that RasGRP1 protein levels were upregulated in the striatum of mice subjected to a cocaine-induced motor sensitization protocol^38^ (data not shown). In addition to diminished ERK and mTORC1 signaling in RasGRP1 KO mice, we also found that pGluR1 (845), a target of protein kinase A (PKA)^39^ was significantly diminished. PKA pathway is reported to be involved in L-DOPA induced dyskinesia^40^. Thus, RasGRP1 can also induce pGluR1 (S845) during L-DOPA-induced dyskinesia. presumably via Ras-cAMP signaling described in yeast^41^.

**Figure 5.**
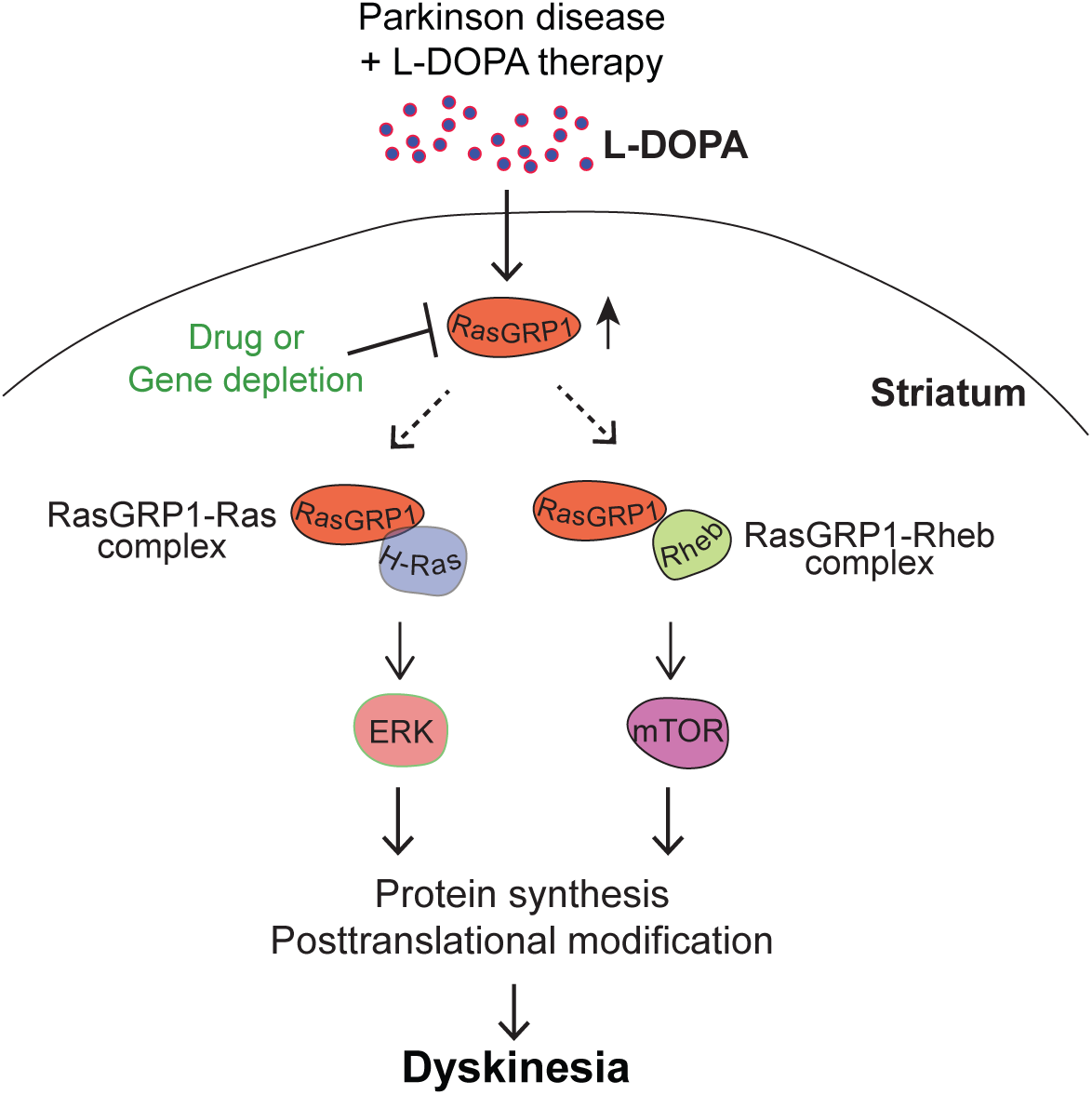
Mechanisms of RasGRP1-induced dyskinesia. Model depicts RasGRP1 is upregulated during L-DOPA-induced dyskinesia and forms complexes with H-Ras to signal ERK, and with Rheb to signal mTOR. Such dual complexes, in parallel, activates ERK and mTOR signaling, exerting a profound cellular and molecular changes in the striatum via protein synthesis and/or protein translation modifications, which will influence the onset and progression of L-DOPA-induced dyskinesia. Drugs or gene depletion strategies that block RasGRP1 may improve the therapeutic efficacy of L-DOPA by diminishing L-DOPA-induced dyskinesia, the debilitating side effects observed in PD patient.

Our proteomics study revealed that RasGRP1 may exert a long-term effect on striatal protein composition to promote strong cellular, molecular, and anatomical alterations via regulating protein synthesis and/or posttranslational modifications. Such biological effects may promote sustained alterations in striatal signaling that could trigger debilitating L-DOPA-induced dyskinesia. A significant number of these putative RasGRP1 proteins could serve as potential targets of L-DOPA-induced dyskinesia. Instead of targeting each of these proteins individually, it may be worth directly targeting RasGRP1, whose deletion is tolerated in mice. Collectively, our study demonstrates that RasGRP1 has a major causal role in L-DOPA induced dyskinesia. Thus, strategies aimed at blocking RasGRP1 may ameliorate L-DOPA-induced dyskinesia in PD patients.

## Acknowledgments

We would like to thank Melissa Benilous for administrative help, and members of the lab for continuous support and collaborative atmosphere. We like to thank members at the animal facility and Scripps proteomics for their help and expertise. This research was supported by grant awards from NIH/NINDS R01-NS087019-01A1, NIH/NINDS R01-NS094577-01A1, and partly from the Cure for Huntington Disease Research Foundation (CHDI).

## Author contributions

S.S conceptualized the project and co-designed the project with A.U. M.E and U. NRJ carried out the lesion and all the behaviors. N.S carried out biochemical and proteomics work. Su.Sw performed cell culture work. N.G helped in Western blotting. O.S purified RasGRP1 and carried out in vitro binding experiments. G. T and C.S.T carried out LC/MS/MS. G. C carried out quantification analysis and bioinformatics. S.S wrote the paper with input from co-authors.

## Data and materials availability

All the data is available in the main text or the supplementary materials.

## Materials and Methods

### Reagents, plasmids, and antibodies

Chemicals and reagents were mainly purchased from Sigma. His-tagged rat CalDAG-GEFII and mouse Flag-CalDAG-GEFI were gift from Ann Graybiel (MIT). The myc-tagged RasGRP1 (pCMV-myc) and GST-tagged (pGEX-6P2) constructs were produced as described^16^. The scrambled shRNA lentiviral control vector was from Addgene, and RasGRP1shRNA was from Sigma (TRCN0000048268-72). Antibodies for RasGRP1 (199), b-Actin, GFAP, GST-HRP, and Myc were obtained from Santa Cruz (sc-8430, sc-47778, sc-33673 and sc-40, respectively). Antibodies against mTOR (2972), pmTOR S2481 (2974), pS6K T389 (9234), pS6 S235/236 (4858), p4EBP1 S65 (9451), p4EBP1 T37/46 (2855), pAkt S473 (4060), pAkt T308 (13038), p44/42 ERK1/2 (9101), S6K (9202), S6 (2217), 4EBP1 (9644), Akt (4691), ERK1/2 (4695), Rheb (13879), pGluR1 S845 (8084), total GluR1 (13185), and DARPP-32 (2306) were from Cell Signaling Technology, Inc. Htt (MAB2166) and TH (MAB318) antibodies were from Millipore-Sigma. Antibodies for GAD1 (10408), GAD2 (21760), PDE1C (13785), PDE2A (55306) were from Proteintech. Rhes antibody (RHES-101AP) was from FabGennix. Glutathione beads were from Amersham Biosciences and Protein G/Protein A agarose beads were obtained from Santa Cruz. Rapamycin, U0126 and wortmannin were from Selleckchem.

### Animals

B6.129P3-*Rasgrp1^tm1Jstn^*/TbwnJ mice (stock #022353) and C57BL/6J mice (stock#000664) were obtained from Jackson Laboratory and maintained in our Scripps animal facility according to IACUC instructions.

### 6-OHDA lesioning

Surgical procedures for unilateral 6-OHDA lesioning were performed as described previously^6^. Briefly, mice were anesthetized deeply by administration of isoflurane. The anesthetized animal was then mounted in a stereotaxic frame (David Kopf Instruments, Tujanga, CA) equipped with a mouse adaptor. 6-OHDA-HCl (Sigma, St Louis, MO) was dissolved in 0.02% ascorbic acid in saline at a final concentration of 2 μg of free-base 6-OHDA per μl. Each mouse received four injections of 6-OHDA (1 μl per injection) into the right striatum, according to the following coordinates(mm): anteroposterior (AP), +1.1; mediolateral (ML), −2; dorsoventral (DV), −3 & −4; and AP +0.1; ML, −2.3; DV, −3.3 & −4.3. Animals were allowed to recover for 3 weeks before behavioral evaluations and L-DOPA treatments were carried out. The efficacy of the lesion was assessed by the loss of tyrosine hydroxylase (TH) in the lesioned striatum (i.e. right) comparing to non-lesioned one (left). Only animals displaying ≥ 60% decrease in striatal tyrosine hydroxylase immunoreactivity were included in statistical analyses of the turning test, drag test, open field, rotarod and AIMs.

### AIMs rating

Abnormal involuntary movements (AIMs) were scored using the rating system described before^6^. Briefly, 6-OHDA-lesioned (or sham) mice were treated for 17 consecutive days with one injection per day of 5 mg/kg L-DOPA plus benserazide (14 mg/kg). AIMs were assessed at day 1, 4, 7, 11, 14, 17 by two observers who were blind to the mice genotypes. On the day of each experiment, mice were habituated in single cages for 30 min, and then received L-DOPA or vehicle injection. 20 min after L-DOPA administration, 3 dyskinetic behaviors were assessed for one min monitoring period which was repeated every 20 min for 120 min. Involuntary movements, distinguished from natural stereotyped behaviors (such as grooming, sniffing, rearing, and gnawing), were classified into three subtypes: locomotive AIMs (tight contralateral turns), axial AIMs (contralateral dystonic posture of the neck and upper body toward the side contralateral to the lesion), limb AIMs (jerky and fluttering movements of the limb contralateral to the side of the lesion). Each subtype was scored on a severity scale from 0 to 4: 0, absent; 1, occasional; 2, frequent; 3, continuous; 4, continuous and not interruptible by outer stimuli. Statistical significance over days was determined by two-way ANOVA (genotype x days of treatment) with repeated measures. Statistical significance over the 120-min test session of each day was determined by two-way ANOVA (genotype x observation periods) with repeated measures, while total AIMs were analyzed by post-hoc comparison.

### Drag test

Mice were habituated in single cages 30 min prior to the test. Each mouse was held 1 cm from the base of the tail and lifted forming an angle of 45 ° from the ground, allowing the support of the front limbs only. The animal was dragged backward and along a surface of 100 cm length at a constant speed of 20 cm/s for five consecutive times (each time alternating the direction of dragging). The animal was video recorded during the whole time. The number of each front limb stepping was counted later by two independent observers watching the videos at a slow pace^42, 43^. Drag test was evaluated 5 days before L-DOPA administration and days 3 and 16 after L-DOPA treatment (1 and 2 hr after injection).

### Open Field

Open field test was used to determinate the total activity. Briefly, animals were placed in the center of each square (50cm x 50cm) open top box under bright light and recorded via ceiling mounted video camera during 40 min. Locomotor activity was assessed using the Ethovision XT11.5 animal tracking software (Noldus) and showed as the total distance travelled in 40 min. Open field test was made 4 days before starting the L-DOPA injections and at fifth day of L-DOPA treatment. In order to quantify the locomotor effect of L-DOPA injection, recording on fifth day of L-DOPA treatment was started after 60 min of the injection.

### Turning test

Recording of turning test was made using the open field boxes. Before beginning the test, one bowl (20 cm diameter) was positioned in the center of the box, then the recording was started after one mouse was placed inside the bowl. The number of contralateral or ipsilateral turns to the 6-OHDA lesion, were automatically detected using the Ethovision XT11.5 animal tracking software (Noldus). Software parameters were adjusted to consider only 360° spin as a quantified turn. Contralateral or ipsilateral turns were quantified during 40 min and expressed as percentage considering total turns (right and left turns) as hundred, only contralateral turns were used for graphs. Turning test was made at 2 days before L-DOPA administration and days 12 of L-DOPA treatment^44, 45^.

### Rotarod

The accelerating rotarod test was used to quantify the motor alterations in wild type and RasGRP1 KO mice. After placing the mice on the rotating rod (diameter of 5 cm), they were tested using the speed of the rotarod accelerated from 4 to 40 RPM over 5-min period and the total time spent on the rod was recorded. In order to identify the hemi-parkinsonian phenotype mice were test on the rotarod after 3 weeks of 6-OHDA lesion and the average of 3 trials was used for analysis. Rotarod evaluation during L-DOPA treatment were made at day 2 and 15, in each day mice were tested using one single trial at 0, 60 and 120 min after L-DOPA injection. In all the cases animals were not trained before the test^7, 46^.

### Cell Culture, Transfections, Amino Acid Treatments and RasGRP1 shRNA experiments

HEK293 cells were cultured in growth medium containing DMEM (Thermo Fischer) with 10% FBS (fetal bovine serum), 1% pen strep, and 5 mM glutamine, essentially as described before^47, 48^. Cells seeded in 3.5- or 6-cm plates, after one day, were transfected with cDNA constructs using polyfect (Qiagen) as per the manufacturer’s instructions. For the amino acid deprivation protocol, after 48 hr, the growth medium was replaced with either serum-free F12+ (D2906, Sigma) medium or F12– (D9785, Sigma) (without L-Leucine) medium for 2 hr, and, wherever indicated, F12– containing cells were stimulated with Leucine (3 mM) medium for the indicated time points, and cells were lysed to proceed with Western blotting. HEK293 cells were cultured in growth medium containing DMEM (Gibco 11965-092) with 10% FBS (fetal bovine serum), 1% pen strep, and 5 mM glutamine as described before^47, 48^ Cells were seeded in 12-well plates for 1 day and then were transfected with RasGRP1 shRNA along with control shRNA constructs using polyfect (Qiagen), according to the manufacturer’s instructions. The growth medium was removed after 24 hr and the cells were lysed to prepare for Western blotting.

### Western blotting, in vivo binding experiments and recombinant protein purification

Protocol for Western blotting, glutathione-binding and Co-IP experiments in striatum was described in our previous studies^16, 47, 49^. GST-RasGRP1 proteins were expressed as described in our earlier work^6, 47, 50^.

### *In vitro* binding

For the *in vitro* binding assay, an equimolar concentration of recombinant purified GST or GST-tagged RasGRP1 were incubated with Rheb for 16 hr at 4°C with glutathione beads in binding buffer containing 50 mM Tris/HCl (pH 8.2), 1 mM dithiothreitol, 100 mM NaCl, and 1% NP-40, and the Rheb was detected through Western blotting with a Rheb antibody, as described earlier^6, 16^.

### TMT labelling, phosphopeptide enrichment and mass spectrometry experimental methods

Striatum from WT and RasGRP1-KO (see scheme Fig 4A) was isolated after L-DOPA treatment were lysed in RIPA buffer precipitated with ice cold acetone overnight and protein resolubilized with 100 μL 6M urea/50 mM TrisHCl, pH 8.0. The protein was subsequently reduced for 45min at 55°C using 3 μL of 0.5 M dithiothreitol (DTT), and then alkylated the dark for 30min using 6 μL of 0.55 M iodoacetamide (IAA). Following reduction and alkylation, protein was once again precipitated with ice cold acetone overnight. The protein pellet was resolubilized with 150 μL TEAB and digestion was performed overnight at 37°C with 6ug trypsin (Promega). Following the digestion, the peptides were quantified using the Pierce^TM^ quantitative colorimetric peptide assay (ThermoFisher Scientific, San Jose, CA). 100 μg of peptides per sample were subsequently labelled with varying TMT labels according to the manufacturer’s instructions (ThermoFisher Scientific, San Jose, CA) and pooled. The pooled plexed samples (400 μg total) were dried under vacuum, resolubilized in 1% TFA and then desalted using OASIS HLB 1cc solid phase extraction cartridges (Waters, Milldford, MA) and then dried once again using vacuum. TMT-labeled phosphopeptides for mass spectrometry were enriched using the High-Select™ Fe-NTA Phosphopeptide Enrichment Kit from ThermoFisher Scientific according to the manufacturer’s instructions. The TMT-labelled non-phoshopeptide complement was cleaned up for mass spectrometry using a C18 ZipTip according to the manufacturer’s instructions (Millipore, Billerica, MA).

For mass spectrometry, dried TMT-labelled peptides (phoshopeptides and non-phosphopeptides) were reconstituted in 5 μL of 0.1% formic acid and on-line eluted into a Fusion Tribrid mass spectrometer (ThermoFisher Scientific, San Jose, CA) from an Acclaim PepMapTM RSLC nano Viper analytical column (75-μm ID × 15 cm, Thermo Scientific, San Jose, CA) using a gradient of 5-25% solvent B (80/20 acetonitrile/water, 0.1% formic acid) in 180 min, followed by 25-44% solvent B in 60 min, 44-80% solvent B in 0.1 min, a 5 min hold of 80% solvent B, a return to 5% solvent B in 0.1 min, and finally a 20 min hold of solvent B. All flow rates were 300 nL/min delivered using a nEasy-LC1000 nano liquid chromatography system (Thermo Fisher Scientific, San Jose, CA). Solvent A consisted of water and 0.1% formic acid. Ions were created at 2.0 kV using the Nanospray FlexTM ion source (ThermoFisher Scientific, San Jose, CA). A synchronous precursor selection (SPS)-MS^3^ mass spectrometry method was used based on the work of Ting et al^51^ scanning between 380-2000 m/z at a resolution of 120,000 for MS1 in the Orbitrap mass analyzer, and performing CID at top speed in the linear ion trap of peptide monoisotopic ions with charge 2-8, using a quadrupole isolation of 0.7 m/z and a CID energy of 35%. The top 10 MS^2^ ions in the ion trap between 400-1200 m/z were then chosen for HCD at 65% energy and detection in the Orbitrap at a resolution of 60,000 and an AGC target of 1E5 and an injection time of 120 msec (MS^3^). Data were analyzed by as described below.

### Proteomic Data Processing and Statistical Analysis

Quantitative analysis of the TMT experiments was performed simultaneously to protein identification using Proteome Discoverer 2.3 software. The precursor and fragment ion mass tolerances were set to 10 ppm, 0.2 Da, respectively), enzyme was Trypsin with a maximum of 2 missed cleavages and Uniprot Mouse proteome FASTA file was used in SEQUEST searches. The impurity correction factors obtained from Thermo Fisher Scientific for each kit was included in the search and quantification. The following settings were used to search the phospho enriched data; dynamic modifications; Oxidation / +15.995Da (M), Deamidated / +0.984 Da (N, Q), Phospho / +79.966 Da (S, T, Y) and static modifications of TMT6plex / +229.163 Da (N-Terminus, K), Carbamidomethyl +57.021 (C). Only unique+ Razor peptides were considered for quantification purposes. Percolator feature of Proteome Discoverer 2.3 was used to set a false discovery rate (FDR) of 0.01. IMP-ptmRS node was used to calculate probability values for each putative phosphorylation site. Total Peptide Abundance normalization method was used to adjust for loading bias and Protein Abundance Based method was used to calculate the protein level ratios. Co-isolation threshold and SPS Mass Matches threshold were set to 50 and 65, respectively. After normal log (In) transformation of the raw ion counts, each channel was divided with reference sample (WT intact). The resulting list of proteins were further filtered by removing proteins that were not quantified in all plexes. Proteins passing this cut off value were exported to JMP (SAS) 13.2.1 for data cleaning and statistical analysis. The non-enriched (aka total) dataset was analyzed in the same fashion except for omission phosphorylation in SEQUEST search and phosphoRS node in PD workflow.

Proteins (including the ones identified by a single peptide) were only included in subsequent analyses if they met the following requirement: the peptides must be quantified in all samples of a given treatment group. Finally, we used one-way ANOVA where Category was fixed effect, to identify proteins that are regulated across comparison groups (Category). The multiple testing correction as per Benjamini Hochberg (B-H) was applied to identify a “top tier” of significant proteins and limit identification of false positives with a False Discovery Rate (FDR) of 10 percent. P values <0.1 were considered statistically significant. The significantly regulated proteins were further interrogated to identify the different treatment group by using Tukey HCD. Proteomic data processing and statistical analysis were carried out as described before^52, 53^.

### Statistical analysis

Data were expressed as mean ± SEM as indicated. All experiments were performed at least in biological triplicate and repeated at least twice. Statistical analysis was performed with a Student’s *t-*test or repeated measure two-way ANOVA followed by post-hoc Bonferroni multiple comparison test or one-way ANOVA followed by Tukey’s multiple comparison test (Graphpad Prism7) as indicated in the figure legends.

---

**Supplementary Figure 1.**
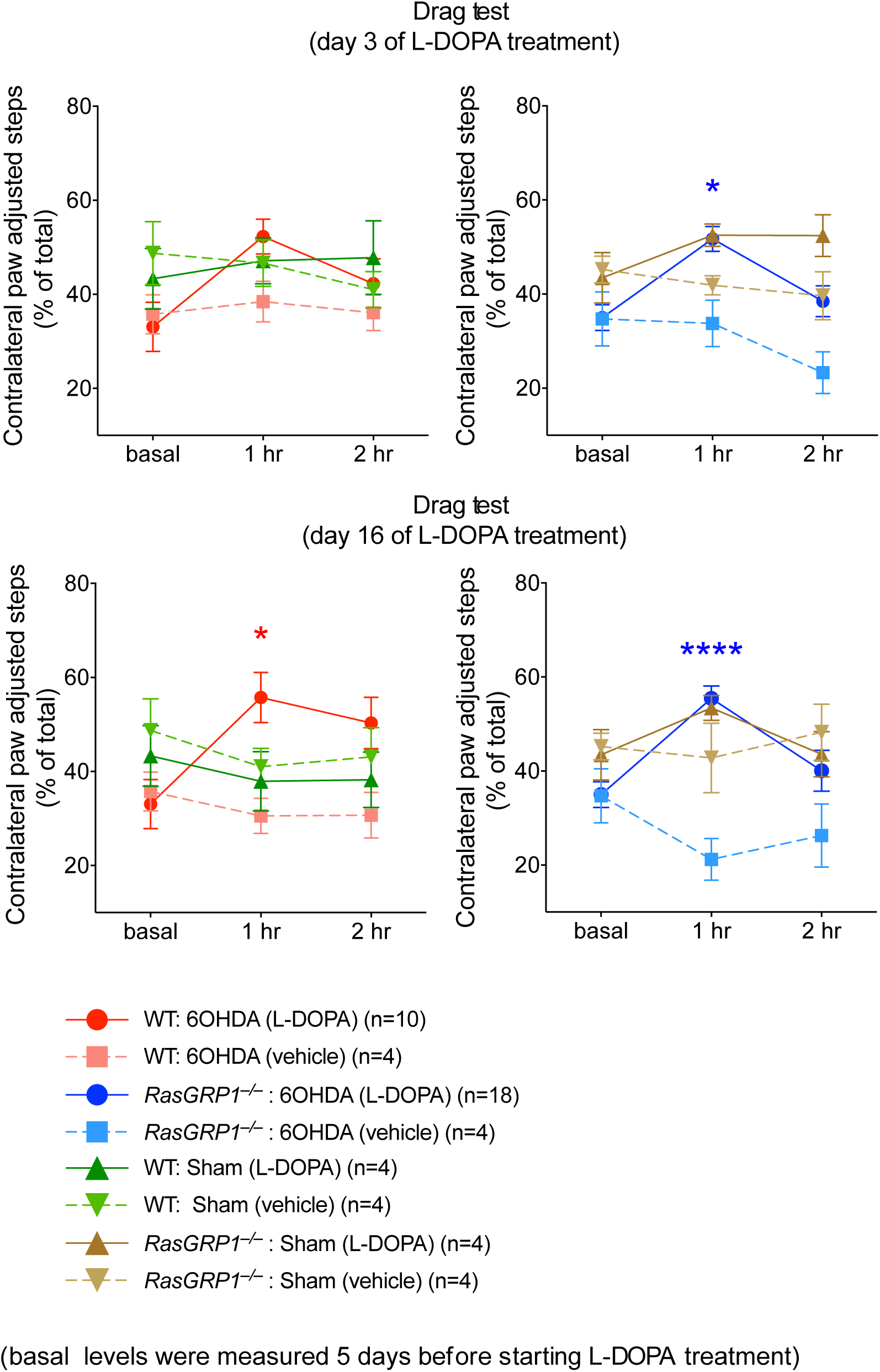
Shows drag test comparing the sham and 6-OHDA lesioned WT and RasGRP1-KO groups on day 3 and day 16 of vehicle or L-DOPA treatment (n = 4-18/group, Two-Way ANOVA followed by Bonferroni post hoc test, *p<0.05; ****p<0.0001).

**Supplementary Figure 2.**
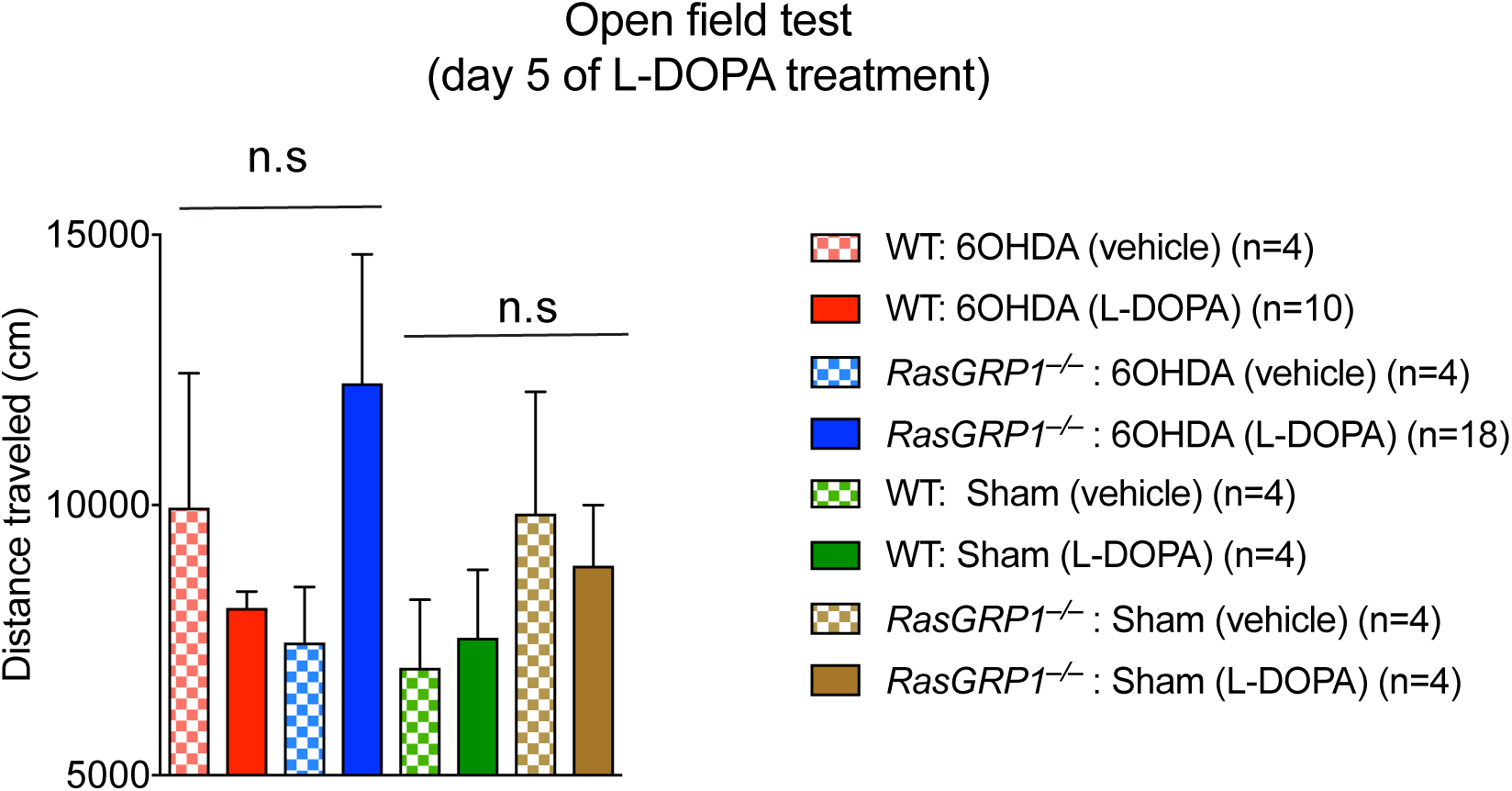
Shows open field test comparing the sham and 6-OHDA lesioned WT and RasGRP1-KO on day 5 of vehicle and L-DOPA treatment. (n = 4-18/group, Two-Way ANOVA followed by Bonferroni post hoc test, n. s; not significant).

**Supplementary Figure 3.**
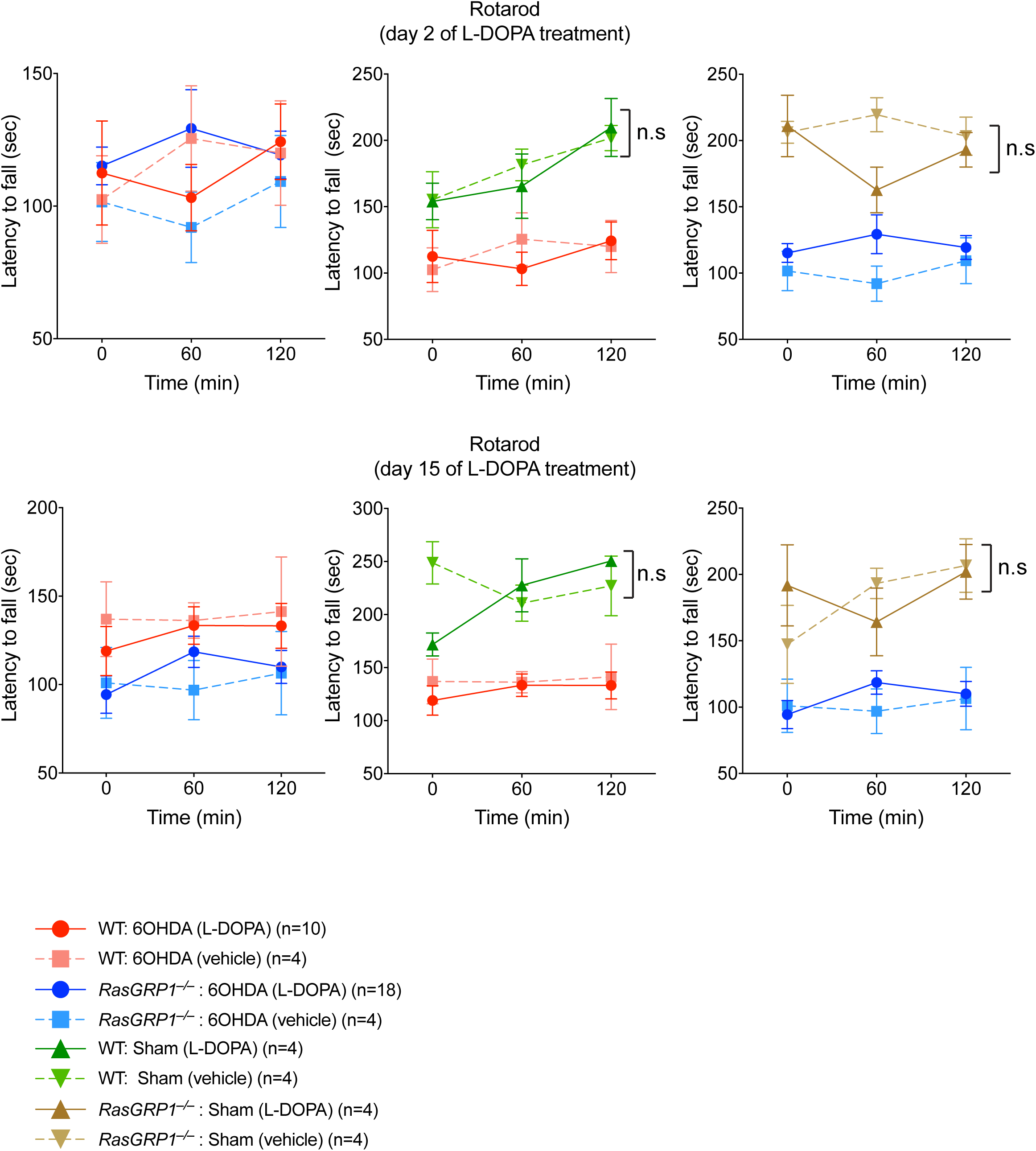
Shows rotarod test after 2 or 15 day of vehicle or L-DOPA treatment in the sham and 6-OHDA lesioned WT and RasGRP1-KO mice (n = 4-18/group, Two-Way ANOVA followed by Bonferroni post hoc test. n. s; not significant).

**Supplementary Figure 4.**
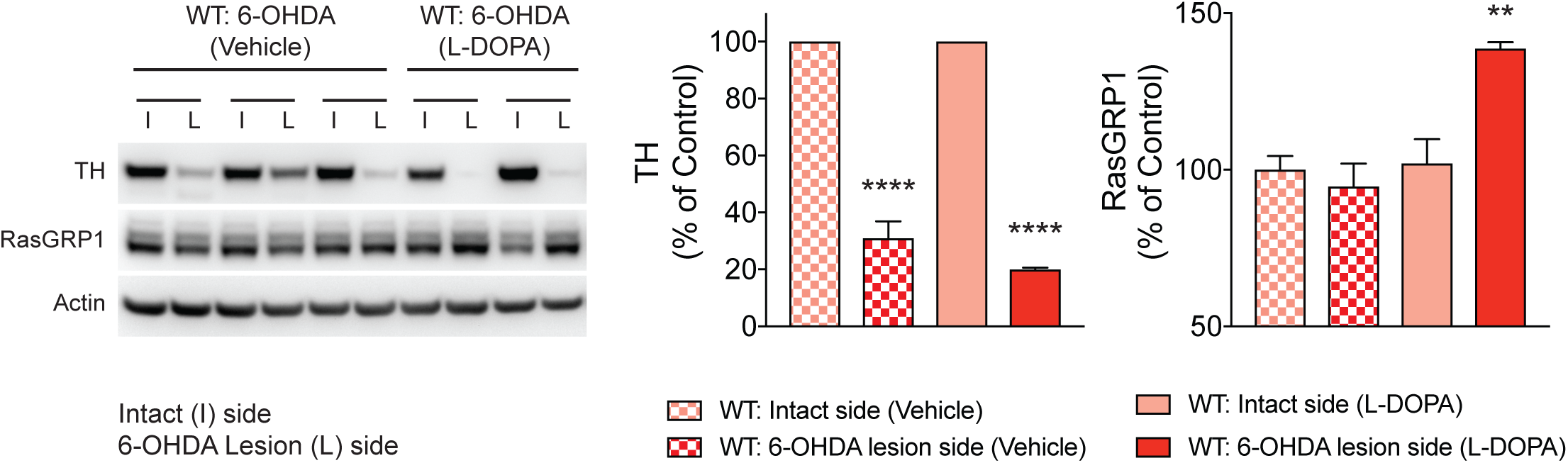
Shows Western blotting of indicated proteins from the striatum of 6-OHDA lesioned WT mice injected with vehicle or L-DOPA. (n = 3/group, One-way ANOVA followed by Tukey’s multiple comparison test, **p<0.01; ****p<0.0001).

**Supplementary Figure 5.**
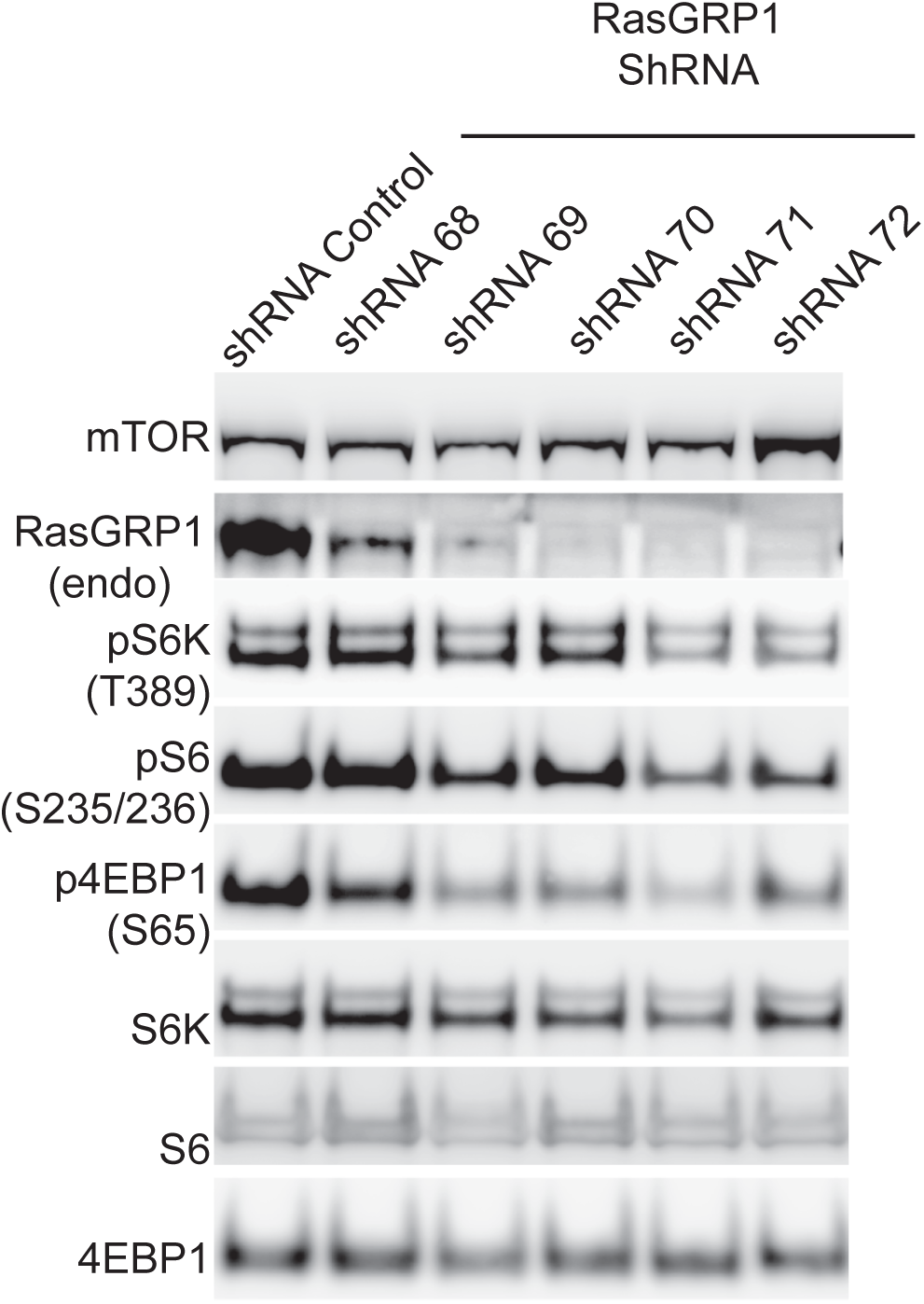
Shows Western blotting of the indicated proteins from HEK293 cells treated with different RasGRP1 shRNA.

